# Treg cells maintain selective access to IL-2 and immune homeostasis despite substantially reduced CD25 function

**DOI:** 10.1101/786228

**Authors:** Erika T. Hayes, Cassidy E. Hagan, Daniel J. Campbell

**Author notes:** Corresponding Author: Daniel J. Campbell, Benaroya Research Institute, 1201 Ninth Avenue, Seattle, WA 98101-2795.

## Abstract

Interleukin-2 (IL-2) is a critical regulator of immune homeostasis through its impact on both regulatory T (Treg) and effector T (Teff) cells. However, the precise role of IL-2 in the maintenance and function of Treg cells in the adult peripheral immune system remains unclear. Here, we report that neutralization of IL-2 abrogated all IL-2 receptor signaling in Treg cells, resulting in rapid dendritic cell (DC) activation and subsequent Teff cell proliferation. By contrast, despite substantially reduced IL-2 sensitivity, Treg cells maintained selective IL-2 signaling and prevented immune dysregulation following treatment with the inhibitory anti-CD25 antibody PC61, even when CD25^hi^ Treg cells were depleted. Thus, despite severely curtailed CD25 expression and function, Treg cells retain selective access to IL-2 that supports their anti-inflammatory functions *in vivo*. Antibody-mediated targeting of CD25 is being actively pursued for treatment of autoimmune disease and preventing allograft rejection, and our findings help inform therapeutic manipulation and design for optimal patient outcomes.

## Introduction

Interleukin-2 (IL-2) is a critical regulator of immune homeostasis through its role in the development, maintenance and function of T regulatory (Treg) cells and its impact on effector cell proliferation and differentiation [1, 2]. The IL-2 receptor (IL-2R) can be composed of 2 or 3 subunits: IL-2R *β* (CD122) and the common gamma (*γ* c) chain (CD132) together form the intermediate affinity receptor, and the addition of IL-2R *α* (CD25) creates the high affinity receptor. Binding of CD25 to IL-2 induces a conformational change that decreases the energy needed to bind to the rest of the receptor, whereas CD122 and CD132 are the critical signaling chains [3]. Treg cells constitutively express CD25, which under homeostatic conditions allows them to outcompete CD25^-^ T effector (Teff) cells and natural killer (NK) cells for limiting amounts of IL-2. This is most important in the secondary lymphoid organs (SLOs), where prosurvival signals downstream of IL-2 signaling maintain Treg cells [4, 5]. Notably, Treg cells cannot make their own IL-2 [6, 7], and therefore depend on IL-2 produced mainly from autoreactive CD4^+^ Teff cells [8, 9]. In this way, Teff and Treg cell populations are dynamically linked and reciprocally control each other to maintain immune homeostasis [10].

When the IL-2-dependent balance of Treg and Teff cells is disrupted, autoimmunity and inflammation can occur. Genetic deficiency in CD25, CD122, or IL-2 results in systemic autoimmune disease in mice [11], and single nucleotide polymorphisms (SNPs) in the *IL2* and *IL2RA* genes are associated with multiple autoimmune diseases in both mice and humans [12, 13]. Therefore, manipulating the IL-2 signaling pathway therapeutically for treatment of autoimmune disease is an area of immense interest. Low dose IL-2 therapy, which enriches Treg cells, has shown efficacy in murine autoimmune models [14–19], and has also benefitted patients with graft versus host disease (GVHD) [20], Hepatitis C virus-induced vasculitis [21], alopecia areata [22], and lupus [23]. However, because IL-2 also acts on effector cells, high dose IL-2 can promote inflammatory responses and this is used for treatment of cancer [24]. As such, safety of therapeutic IL-2 remains a concern, and efficacy can vary widely depending on the current disease activity and immune history of the patient. Indeed, in two mouse models of type 1 diabetes, early intervention with IL-2 prevented disease, but initiation of treatment after loss of tolerance (but before overt hyperglycemia) accelerated disease progression [13, 19]. The fact that monoclonal antibodies against CD25 are also used as an immunosuppressive to treat organ transplant rejection [25] and demonstrated efficacy against multiple sclerosis (MS) [26] further highlights the complexity of targeting this signaling pathway.

The inhibitory anti-CD25 antibody PC61 has been extensively used to examine the role of CD25 in IL-2 signaling in Treg cells in mice [27, 28], and model the impact of blocking IL-2 signaling *in vivo*. However, interpretation of results is difficult due to uncertainty of whether the observed *in vivo* effects are mediated by functional blockade of CD25, Treg cell depletion, or a combination [29–31]. Using PC61 derivatives with identical epitope specificity but divergent constant region effector function, a recent study showed that only depletion of CD25^hi^ cells and not blockade of CD25 could disrupt immune homeostasis [32]. However, the fact that blockade of CD25 for up to four weeks caused no disturbance in immune homeostasis is surprising, given the central role IL-2 is thought to play in the maintenance of Treg cells in SLOs. For instance, acute blockade of IL-2 using the IL-2 antibody S4B6-1 (S4B6) significantly reduces Treg cells, and when administered early in life causes Treg cell dysfunction sufficient to induce autoimmune gastritis in Balb/c mice [8]. However, in addition to blocking IL-2 binding to CD25, this antibody forms superagonistic IL-2 immune complexes that are specifically targeted to CD122^hi^ effector populations such as NK cells and memory T cells and this may have contributed to disease development in these animals. These divergent results may reflect differences in the importance of IL-2 for the induction vs. maintenance of immune tolerance, or may reflect idiosyncrasies in how the reagents used for IL-2 and CD25 blockade actually impact IL-2 availability and signaling in Treg and Teff cells.

In light of this confusion, we comprehensively examined how manipulating the IL-2/CD25 axis by different methods perturbs Treg cell maintenance, phenotype and function in maintaining normal immune homeostasis. We found that neutralization of IL-2 abrogated all STAT5 phosphorylation (pSTAT5) in Treg cells, resulting in rapid dendritic cell (DC) activation and subsequent Teff cell proliferation and expansion. By contrast, Treg cells maintained normal IL-2 signaling in the presence of the inhibitory anti-CD25 antibody PC61 *in vivo*, despite substantially reduced sensitivity to IL-2. Continued IL-2 signaling was dependent on residual CD25 function, and we found that even CD25^lo^ Treg cells that escape depletion after treatment with a strongly depleting IgG2a version of PC61 initially maintain IL-2 responsiveness and functionality *in vivo*. However, prolonged anti-CD25-mediated Treg cell depletion results in loss of immune homeostasis and Teff cell proliferation and activation. These findings demonstrate that even with severely curtailed CD25 function, Treg cells retain their selective access to IL-2 *in vivo*, and this is sufficient to maintain normal Treg cell function and immune homeostasis. These data warrant re-examination of previous studies using the PC61 antibody [31, 32], and have important implications for efforts to target the IL-2/CD25 axis therapeutically to dampen inflammation and induce immune tolerance.

## Results

### Complete antibody-mediated neutralization of IL-2 in vivo

Following administration *in vivo*, the anti-IL-2 monoclonal antibodies S4B6 and JES6-1A12 (JES6) can complex with endogenous IL-2 and act as super-agonists for different leukocyte populations, depending on the antibody used and IL-2R component expression of the cell [33]. However, co-administration of S4B6 and JES6 should block binding to both CD122 and CD25, and effectively neutralize all IL-2 function. To test this, we treated mice with either S4B6 alone, equal amounts of S4B6 and JES6, or an excess of JES6 over S4B6, and assessed IL-2 signaling and activation of Treg cells and NK cells after seven days. Due to the qualitative difference in response to IL-2 in Treg cells (bimodal) compared to NK cells (a weaker unimodal shift) (Fig. S1A), we reported response to IL-2 as frequency pSTAT5^+^ of Treg cells and geometric mean fluorescence intensity (gMFI) of NK cells, respectively. S4B6 blocks IL-2 binding to CD25, and targets IL-2 to cells expressing high levels of CD122, and accordingly treatment with S4B6 alone induced robust proliferation of NK cells associated with increased STAT5 phosphorylation (Fig. 1A). However, the proliferation of NK cells as well as their elevated STAT5 phosphorylation was completely blocked by the addition of an equal amount of the JES6 antibody that inhibits IL-2/CD122 binding. As NK cells express very high levels of CD122 and are potently stimulated by S4B6/IL-2 immune complexes, these data demonstrate addition of JES6 prevents the formation of super-agonistic IL-2/S4B6 immune complexes *in vivo*. Furthermore, consistent with the ability of S4B6 to effectively block IL-2 interaction with CD25, all treatments potently inhibited STAT5 phosphorylation in Treg cells (Fig. 1B). Thus, this antibody combination can completely neutralize IL-2 activity on multiple cell types *in vivo*, and out of an abundance of caution we used an excess of JES6 over S4B6 for the anti-IL-2 treatment in all subsequent experiments.

**Figure 1.**
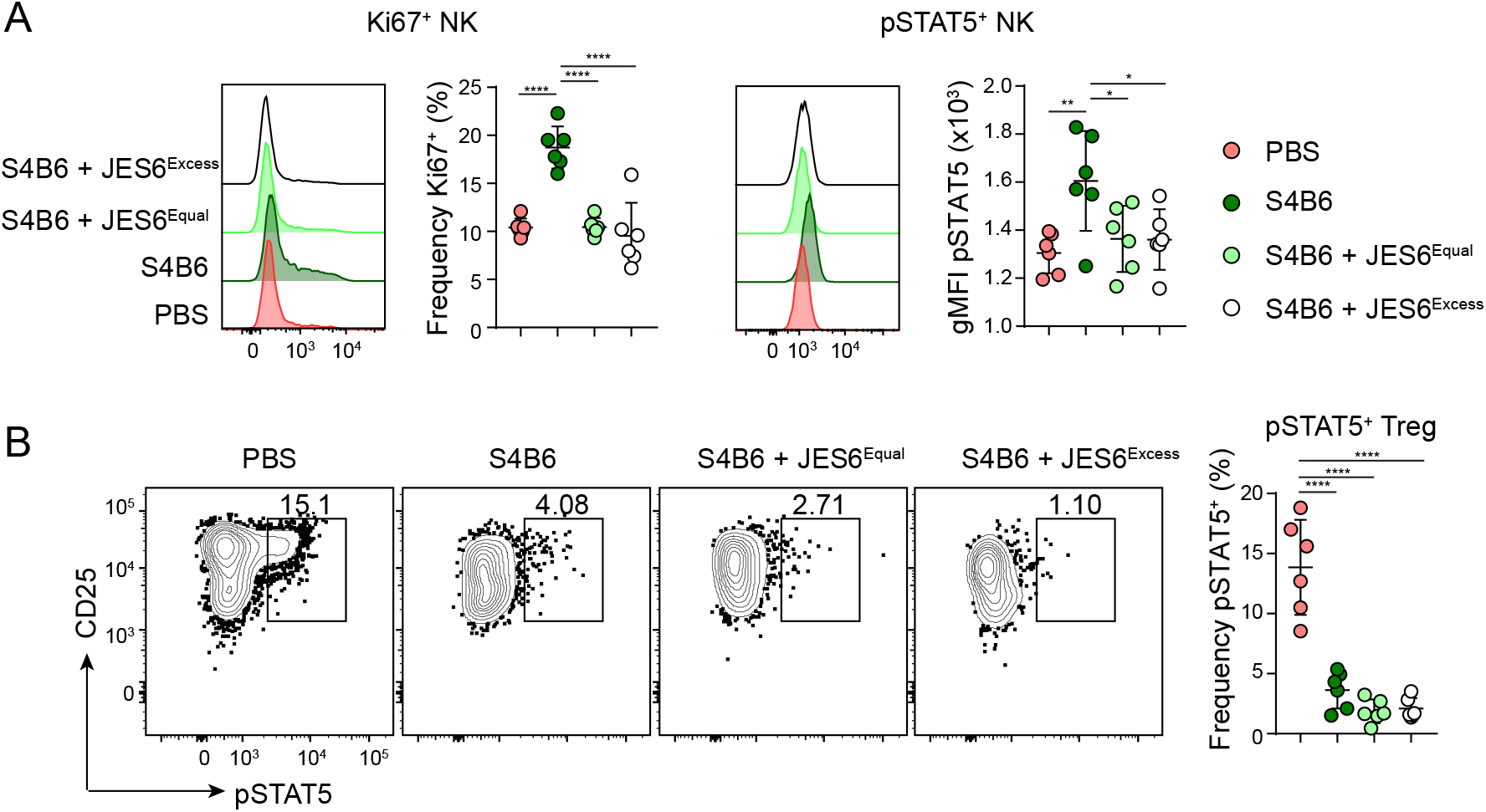
The combination of JES6 and S4B6 effectively neutralizes IL-2 *in vivo*. WT B6 mice were treated IP with PBS, 150μg S4B6 alone (no JES6), 150μg S4B6 and 150μg JES6, or 500μg JES6 and 150μg S4B6 on day 0 and day 5, and sacrificed on day 7 for analysis. (A) Representative flow cytometric analyses of Ki67 (left) and pSTAT5 (right) expression in gated NK1.1^+^ splenic NK cells in each treatment group. Corresponding graphical analysis of frequency of Ki67^+^ and gMFI of pSTAT5 in splenic NK cells. (B) Representative flow cytometry analysis of pSTAT5 and CD25 expression by gated splenic Foxp3^+^ Treg cells. Right, graphical analysis of frequency of pSTAT5^+^ Treg cells in each treatment group. Data is combined from two independent experiments, 6 mice per group total. Significance determined by one-way ANOVA with Tukey post-test for pairwise comparisons. *p<0.05, **p<0.01, ***p<0.001, ****p<0.0001.

### Differential impacts of targeting CD25 or IL-2 on CD25^+^ T cells

To compare how inhibiting the IL-2/CD25 axis by targeting either CD25 or IL-2 impacts Treg cell abundance and immune homeostasis, C57BL/6 (B6) mice were treated intraperitoneally (IP) with an engineered isoform of PC61 (PC61^N297Q^) that inhibits CD25 function but does not deplete CD25-expressing cells, an engineered isoform of PC61 (PC61^2a^) that has strong depleting activity, or a combination of S4B6 and JES6 as above. In line with previously reported findings [32], seven days after treatment Treg cells were reduced by ~50% in mice treated with either PC61^2a^ or anti-IL-2 relative to PBS-treated controls (Fig. 2A). Surprisingly, no significant change in the frequency or absolute number of Treg cells was observed in PC61^N297Q^ treated mice. Treg cells can be divided into central (c)Treg and effector (e)Treg cells based on differential expression of CD62L and CD44. In both PC61^2a^-and anti-IL-2-treated mice, there was a specific loss of CD62L^+^CD44^lo^ cTreg cells (Fig. 2B), which express the highest levels of CD25 and are the most dependent on IL-2 for their homeostatic maintenance within the spleen [34]. Staining isolated cells with a flourochrome conjugated PC61 seven days after PC61^N297Q^ and PC61^2a^ treatment showed essentially complete coverage of the epitope (Fig. 2A and Fig. 2C), verifying that we used a saturating concentration of injected antibody. To assess CD25 expression in treated animals, we stained cells with the 7D4 anti-CD25 antibody, which binds a distinct epitope and does not compete with PC61 for binding. As expected, PC61^2a^ treatment effectively depleted CD25^hi^ cells, and the remaining Treg cells in these animals were CD25^mid/lo^. Similarly, CD25 expression was significantly reduced in anti-IL-2-treated mice, which is likely due to the ability of IL-2 signaling and activated STAT5 to promote CD25 expression in a positive feedback loop [35]. Interestingly, despite lacking the ability to deplete CD25^+^ cells, CD25 expression was also significantly decreased on Treg cells from mice that had been treated with PC61^N297Q^ (Fig. 2C), indicating that this antibody may induce surface cleavage or internalization of CD25. Finally, CD4^+^ and CD8^+^ Teff cells can transiently express high levels of CD25 upon activation, and thus could also be affected by the treatments administered. While very few CD8^+^ Teff cells expressed CD25 in any treatment group (not shown), about 2% of Foxp3^-^CD44^+^ CD4^+^ Teff cells were CD25^+^ (Fig. 2D) and both PC61^N297Q^ and PC61^2a^ treatment significantly reduced the frequencies and absolute numbers of this population.

**Figure 2.**
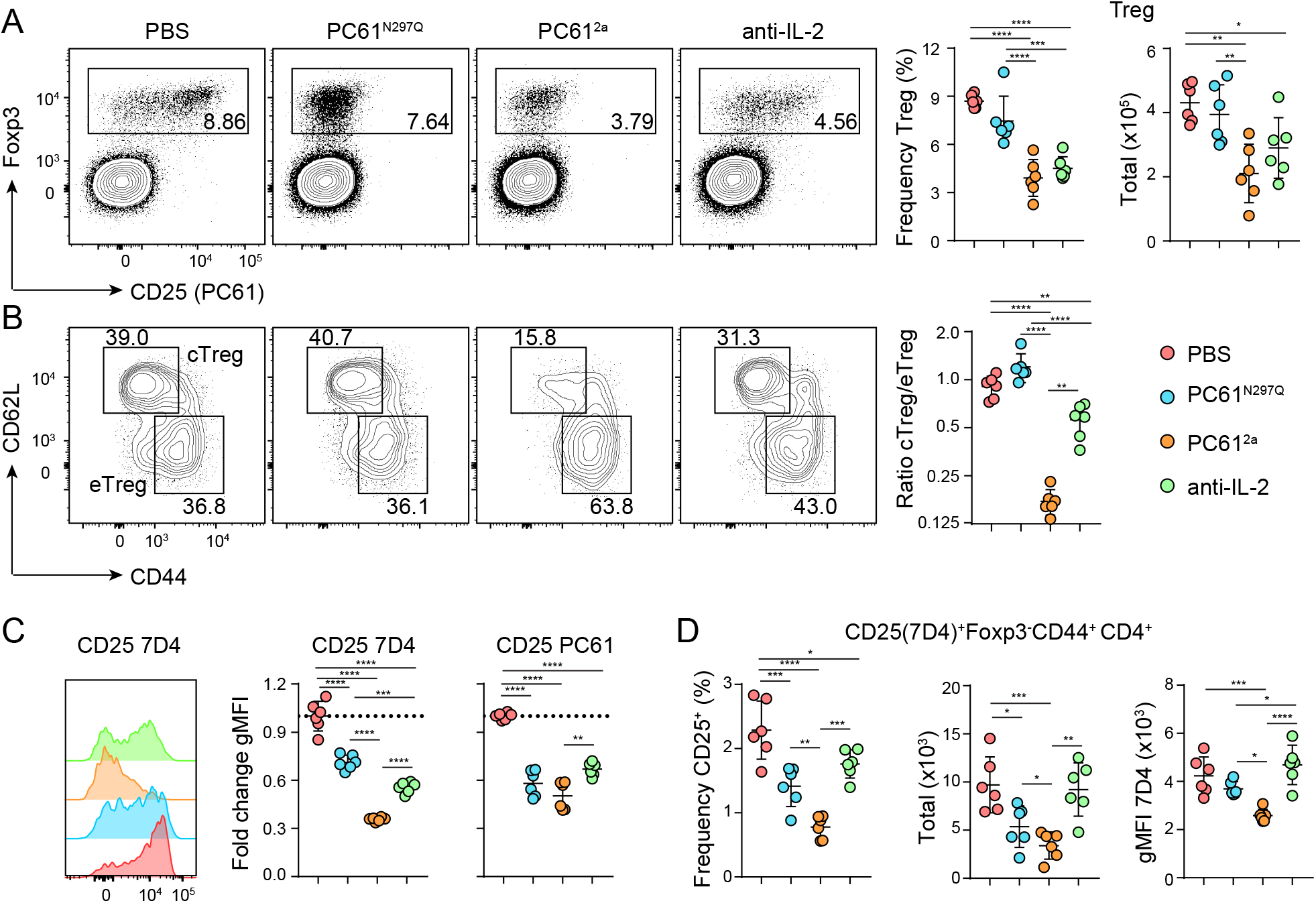
Impacts of targeting CD25 or IL-2 on Treg cells. WT B6 mice were treated IP with PBS, PC61^N297Q^, PC61^2a^, or anti-IL-2 (S4B6 + JES6), and sacrificed for analysis after seven days. (A) Representative flow cytometric analysis of Foxp3 and CD25 (clone PC61) expression by gated splenic CD4^+^ T cells. Foxp3^+^ Treg cells are gated as indicated. Right, Graphical analysis of frequency and total number of splenic Treg cells in each treatment group. (B) Representative flow cytometry analysis of CD44 and CD62L expression by gated splenic Foxp3^+^ Treg cells showing gates used to define cTreg and eTreg populations. Right, graphical analysis of the ratio of cTreg cells to eTreg cells in the spleens of each treatment group. (C) Representative flow cytometry histograms of CD25 7D4 staining in Treg cells. Right, graphical analysis of fold change in gMFI over controls of CD25 PC61 and CD25 7D4 staining by Treg cells in each treatment group. (D) Graphical analysis of frequency, total number, and gMFI of CD25^+^ (7D4) Foxp3^-^CD44^+^CD4^+^ splenic T cells in each treatment group. Data is combined from two independent experiments, 6 mice per group total. Significance determined by one-way ANOVA with Tukey post-test for pairwise comparisons. *p<0.05, **p<0.01, ***p<0.001, ****p<0.0001.

### IL-2 neutralization leads to cDC2 activation and drives autoreactive Teff cell proliferation

A critical function of Treg cells in SLOs is to restrain DC activation and prevent excessive T cell priming. Therefore, to assess the functionality of Treg cells in the anti-CD25 and anti-IL-2 treated mice, we examined the DC abundance and activation status in the spleen by measuring expression of CD86 and CD40, two important costimulatory molecules that are upregulated in activated DCs. Although no changes were observed in 33D1^-^CD11b^-^ type-1 conventional DCs (cDC1) or 33D1^-^CD11b^hi^ monocyte-derived DCs (moDCs) (Fig. S1B), expression of CD86 and CD40 was elevated on 33D1^+^CD11b^+^ type-2 conventional DCs (cDC2) in anti-IL-2 treated mice (Fig. 3A). Along with the increase in activated DCs, the anti-IL-2 treated mice also showed enhanced proliferation of the Foxp3^-^CD44^+^ CD4^+^ and CD44^+^CD62L^+^ CD8^+^ Teff cell populations (Fig. 3B). No significant changes in proliferating NK cells were observed, nor in any cell type in mice treated with PC61^N297Q^ or PC61^2a^ compared to controls. Thus, although both IL-2 neutralization and PC61^2a^ treatments resulted in a similar numeric loss of Treg cells, increased DC activation and proliferation of Teff cells was only observed during IL-2 blockade.

**Figure 3.**
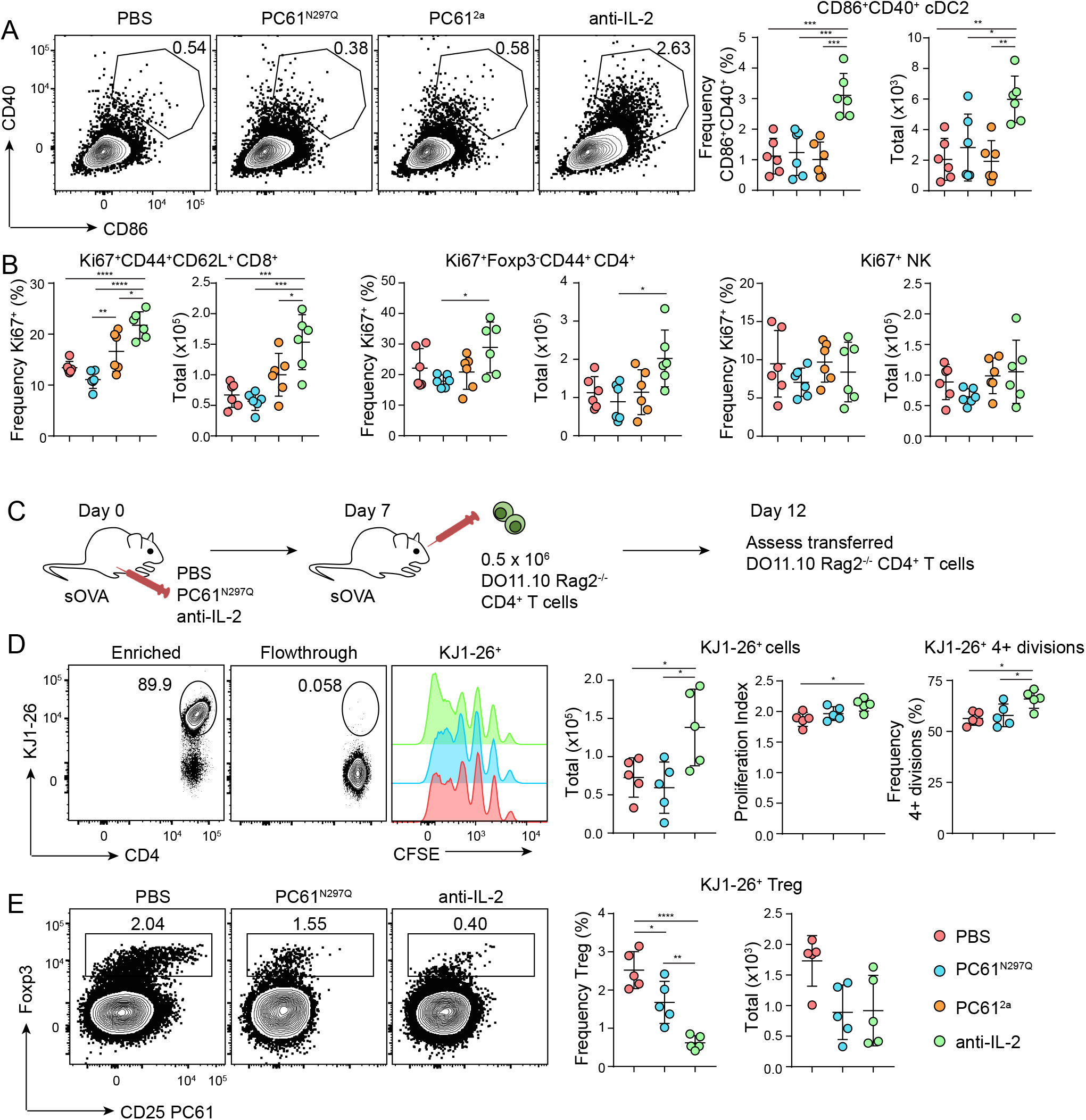
IL-2 neutralization leads to cDC2 activation and drives autoreactive Teff cell proliferation. (A-B) WT B6 mice were treated IP with PBS, PC61^N297Q^, PC61^2a^, or anti-IL-2 (S4B6 + JES6), and sacrificed after seven days for analysis. (A) Representative flow cytometric analysis of CD86 and CD40 expression by gated splenic MHCII^+^CD11c^+^33D1^+^CD11b^+^ cDC2. Right, graphical analysis of frequency and total number of splenic CD86^+^CD40^+^ cDC2s in each treatment group. (B) Graphical analysis of frequency and total number of splenic Ki67^+^CD44^+^CD62L^+^ CD8^+^ T cells, Ki67^+^Foxp3^-^CD44^+^CD4^+^ T cells, and Ki67^+^ NK cells in each treatment group. (C-E) Balb/c.sOVA mice were treated IP with PBS, PC61^N297Q^ or aIL-2. After seven days, 0.5×10^6^ DO11.10 Rag2^-/-^ CD4^+^ T cells were transferred retro-orbitally into each mouse, and animals were sacrificed five days later for analysis. (C) Experimental schematic. (D) Representative flow cytometric analysis of the KJ1-26 DO11.10 clonotype on transferred CD4^+^ T cells on an enriched spleen sample versus flowthrough. Right, representative histograms of cell proliferation based on CFSE dilution in gated KJ1-26^+^ T cells from each treatment group. Graphical analyses of total splenic cells recovered, proliferation index, and frequency of cells having undergone four or more divisions in transferred KJ1-26^+^ cells in each treatment group. (E) Representative flow cytometric analysis of Foxp3 and CD25 (PC61) expression by transferred KJ1-26^+^ cells. Foxp3^+^ Treg cells are gated as indicated. Right, Graphical analysis of frequency and total number of splenic KJ1-26^+^ Treg cells in each treatment group. Data is combined from two independent experiments, 5-6 mice per group total. Significance determined by one-way ANOVA with Tukey post-test for pairwise comparisons. *p<0.05, **p<0.01, ***p<0.001, ****p<0.0001.

We hypothesized that impaired Treg cell function with IL-2 neutralization led to the increased costimulatory molecule expression on cDC2s, which in turn allowed these DCs to more potently activate naïve autoreactive T cells. To directly test this, we used mice expressing a soluble form of ovalbumin (sOVA) that is efficiently processed and presented on the surface of splenic cDC2s [9, 36]. sOVA mice were treated with PBS, PC61^N297Q^, or anti-IL-2, and on day 7 given 0.5×10^6^ CFSE-labelled naïve CD4^+^ OVA-specific T cells from DO11.10 Rag2^-/-^ mice. Five days post transfer we assessed the proliferation and activation of the DO11.10 cells (identified by staining with the clonotype-specific antibody KJ1-26) in the spleen following their magnetic enrichment (Fig. 3C, D). Although transferred antigen-specific T cells proliferated in all recipient mice, the total number of transferred cells recovered, the proliferation index, and the frequency of cells that had 4 or more divisions were all significantly higher in the anti-IL-2 treated mice (Fig. 3D), indicating that altered cDC2 activation following IL-2 blockade promotes the enhanced activation and proliferation of autoreactive CD4^+^ T cells. We also assessed the ability of peripheral (p)Treg cells to develop from the transferred naïve T cells. Indeed, the frequency and number of splenic Foxp3^+^ DO11.10 T cells was significantly reduced by both PC61^N297Q^ and anti-IL-2 treatments (Fig. 3E). Host cells from recipient mice showed the same phenotypes as described in Figures 2 and 3 (Fig. S1C) in response to PC61^N297Q^ and anti-IL-2 treatment. Collectively, these data confirm the critical role that IL-2 plays in maintaining Treg-dependent DC homeostasis and supporting pTreg cell differentiation in the lymphoid organs, and demonstrate that acute IL-2 neutralization leads to excessive T cell priming and activation.

### Treg cells retain selective access to IL-2 in a CD25-dependent manner in the presence of PC61

The lack of any immune dysregulation in mice treated with PC61^N297Q^ or PC61^2a^ suggests that Treg cells in these animals retain significant functional capacity compared with Treg cells in anti-IL-2 treated mice. Indeed, differences in IL-2 signaling in cells subject to these treatments could explain the alterations observed in Treg cell number and function, as well as immune activation status. Therefore, we examined pSTAT5 directly *ex vivo* in different cell populations one week after antibody administration, when PC61 epitopes on CD25 were still completely saturated (Fig. 2A, C). Whereas neutralization of IL-2 blocked all pSTAT5 as expected, Treg cells from animals treated with PC61^N297Q^ maintained normal levels of pSTAT5, and even treatment with PC61^2a^ had only a modest impact on the frequency of pSTAT5^+^ Treg cells (Fig. 4A). The pSTAT5 staining we observed in the treated animals does not simply reflect prolonged IL-2 signaling that occurred prior to treatment initiation, as we have previously shown that injection of IL-2 antibodies as little as 30 minutes prior to sacrifice ameliorates all detectable pSTAT5 in Treg cells [9]. Treatment with PC61^N297Q^ or PC61^2a^ also did not redirect IL-2 to effector cells, as the gMFI of pSTAT5 in both NK cells and CD44^+^CD62L^+^ CD8^+^ Teff was not increased by any of the treatments (Fig. S2A). Thus, although numbers of Treg cells are similarly decreased in the PC61^2a^ and anti-IL-2-treated animals, sustained IL-2 signaling in PC61^2a^-treated mice likely helps maintain Treg cell function and prevents the overt immune dysregulation that occurs in anti-IL-2-treated animals. IL-2 signaling helps maintain Treg cell function by promoting high expression of Foxp3 [37] and other inhibitory molecules like CTLA-4 [38]. Consistent with this, we found that the gMFI of Foxp3 was significantly reduced in anti-IL-2 treated mice, whereas PC61^N297Q^ and PC61^2a^ had only minor impacts on Foxp3 expression (Fig. 4B). Furthermore, Treg cells from PC61^2a^ treated mice had elevated expression of CTLA-4 compared to anti-IL-2 treated mice (Fig. 4C).

**Figure 4.**
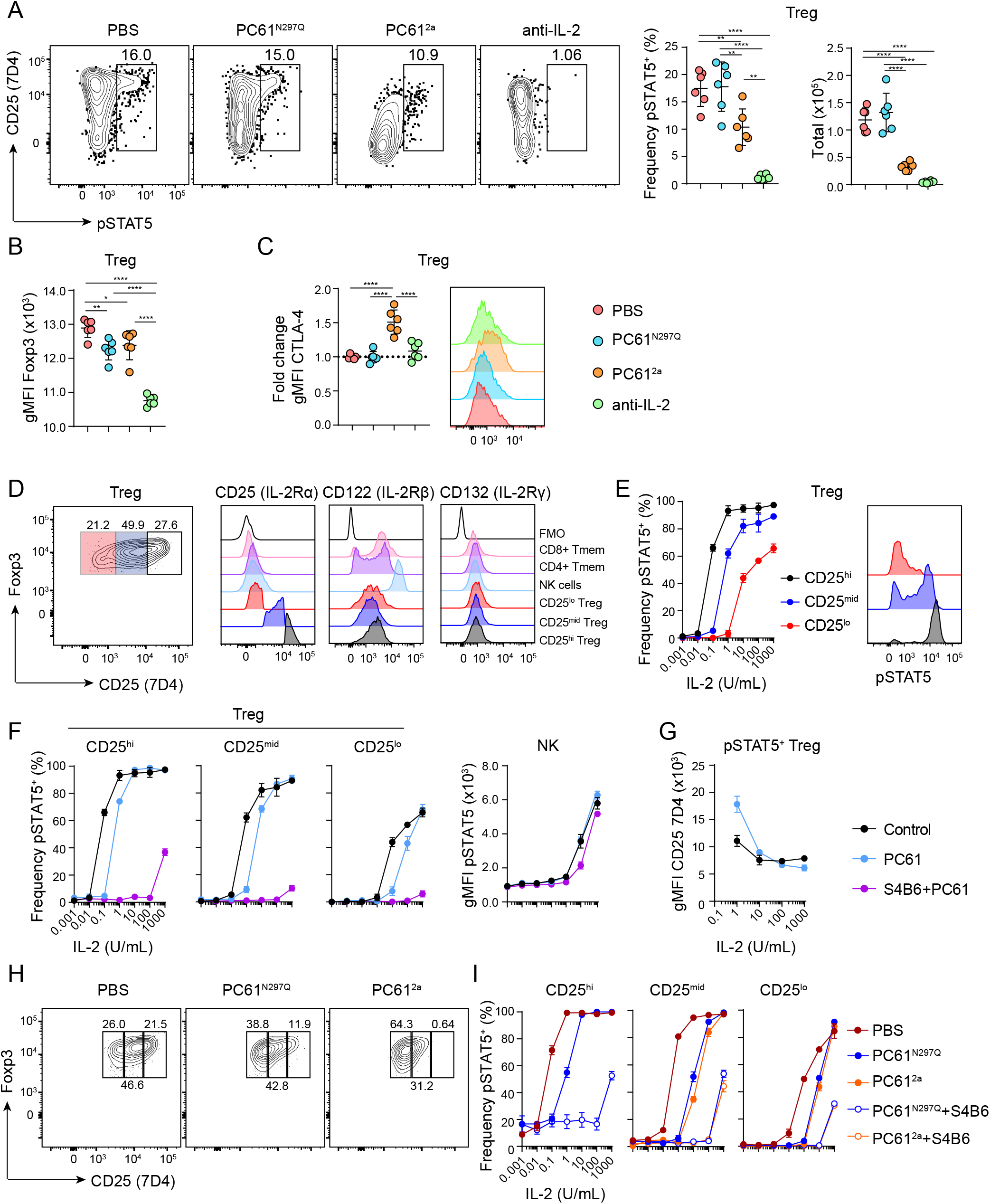
Treg cells retain selective access to IL-2 *in vivo* despite severely curtained CD25 function. (A-C) WT B6 mice were treated IP with PBS, PC61^N297Q^, PC61^2a^, or anti-IL-2 (S4B6 + JES6), and sacrificed after seven days for analysis. (A) Representative flow cytometric analysis of pSTAT5 and CD25 (7D4) by gated splenic Treg cells. Right, graphical analysis of frequency and total number of pSTAT5^+^ splenic Treg cells in each treatment group. (B) Graphical analysis of gMFI Foxp3 in gated Foxp3^+^ Treg cells in the spleens of each treatment group. (C) Graphical analysis of fold change in gMFI over controls of CTLA-4 staining by Treg cells in each treatment group. Right, representative flow cytometric analysis of CTLA-4 by gated splenic Treg cells. (D) Representative flow cytometric analysis of CD25 (7D4) expression in untreated Treg cells, and gates defining CD25^hi^, CD25^mid^ and CD25^lo^ cells are shown. Right, representative flow cytometric analysis of CD25, CD122 and CD132 expression by the indicated cell populations and fluorescence minus one (FMO) controls. (E) Graphical analysis of frequency of pSTAT5^+^ splenic Treg cells within each CD25 expression subset in response to rIL-2. Right, representative flow cytometric analysis of pSTAT5 staining in Treg cells in each CD25 subset in response to 1 U/mL rIL-2. (F) Graphical analysis of frequency of pSTAT5^+^ Treg cells in response to IL-2 after treatment *in vitro* with either PC61, or PC61 and S4B6 compared to controls. Right, graphical analysis of gMFI of pSTAT5 in NK cells under the same treatment conditions. (G) Graphical analysis of gMFI of CD25 (7D4) in pSTAT5^+^ Treg cells in response to IL-2 after treatment *in vitro* with PC61 compared to controls. (H-I) WT B6 mice were treated IP with PBS, PC61^N297Q^ or PC61^2a^ and sacrificed after 24 hours. (H) Representative flow cytometry analysis of CD25 (7D4) staining on gated Treg cells, and gates defining CD25^hi^, CD25^mid^ and CD25^lo^ cells are shown. (I) Graphical analysis of frequency of pSTAT5^+^ Treg cells in response to *in vitro* IL-2 +/-S4B6 in each *in vivo* treatment group (PBS, PC61^N297Q^ or PC61^2a^) as indicated. (A-C) Data is combined from two independent experiments, 6 mice per group total. Significance determined by one-way ANOVA with Tukey post-test for pairwise comparisons. *p<0.05, **p<0.01, ***p<0.001, ****p<0.0001. (D-I) Data is from one representative experiment, with three technical replicates per condition. Experiments were repeated independently at least three times.

The ability of Treg cells to maintain IL-2 responsiveness in the presence of the PC61 antibodies led us to two competing hypotheses. Either the CD25 remaining on the cell surface was still functional and mediating IL-2 signaling, or residual IL-2 signaling was CD25-independent, perhaps due to upregulation or increased sensitivity of the other IL-2R components on Treg cells, or due to changes in the IL-2R signaling pathways such as downregulation of protein phosphatase 2A (PP2A) [39]. Co-staining with the 7D4 anti-CD25 antibody clearly showed that as in control mice, pSTAT5 was enriched among Treg cells expressing the highest amounts of CD25 in both PC61^N297Q^-and PC61^2a^-treated mice (Fig. 4A). We therefore compared the expression of the other IL-2R components from untreated splenic Treg cells divided into three subsets based on their expression of CD25 by 7D4 staining. Expression of CD122 and CD132 was similar between all three subsets of Treg cells. Furthermore, CD122 expression by Treg cells was much lower than expression by memory T cells or NK cells. Thus, enhanced expression of the intermediate affinity IL-2R cannot explain the ability of CD25^hi^ Treg cells to selectively respond to IL-2 in the presence of the PC61 antibodies (Fig. 4D).

To directly assess the effect that the PC61 has on CD25 function and the sensitivity of splenic Treg cells to IL-2, we performed *in vitro* stimulations in the presence of PC61 and the anti-IL-2 clone S4B6, which directly blocks interaction between IL-2 and CD25 but has minimal impact on IL-2 signaling via the intermediate affinity CD122/CD132 complex [40]. For analysis, Treg cells were subsetted based on their expression of CD25 by 7D4 staining as in Fig 4D. CD25^hi^ Treg cells achieved maximal pSTAT5 at a relatively low dose of rIL-2 (1 U/mL), while CD25^mid^ Treg cells were approximately 10-fold less sensitive and CD25^lo^ Treg cells were more than 100-fold less sensitive (Fig. 4E). Pre-treatment with PC61 for 30 min prior to IL-2 stimulation reduced IL-2 sensitivity by ~10-fold in all three Treg cell populations (Fig 4F), but all were still able to achieve the maximal level of pSTAT5 observed in untreated cells. However, further addition of S4B6 severely curtailed IL-2 sensitivity in all Treg cells, indicating that they do not efficiently signal through the intermediate-affinity IL-2 receptor. By contrast, in NK cells, which lack CD25 but have high levels of CD122, IL-2 responses were completely unaffected by the addition of PC61 (Fig. 4F, right), and S4B6 had only a small effect on signaling which is due to minor steric inhibition *in vitro* [40]. Thus, we conclude that CD25 retains significant functionality when bound by PC61. Indeed, PC61 does not directly occlude IL-2 binding, but rather inhibits CD25 function by inducing a conformational change in the IL-2 binding pocket [41]. In fact, at lower doses of IL-2 (1 U/mL), treatment with PC61 actually makes Treg cells more CD25 dependent, as evidenced by the increased CD25 gMFI of pSTAT5^+^ Treg cells treated with PC61 compared to untreated control cells (Fig. 4G).

**Figure 5.**
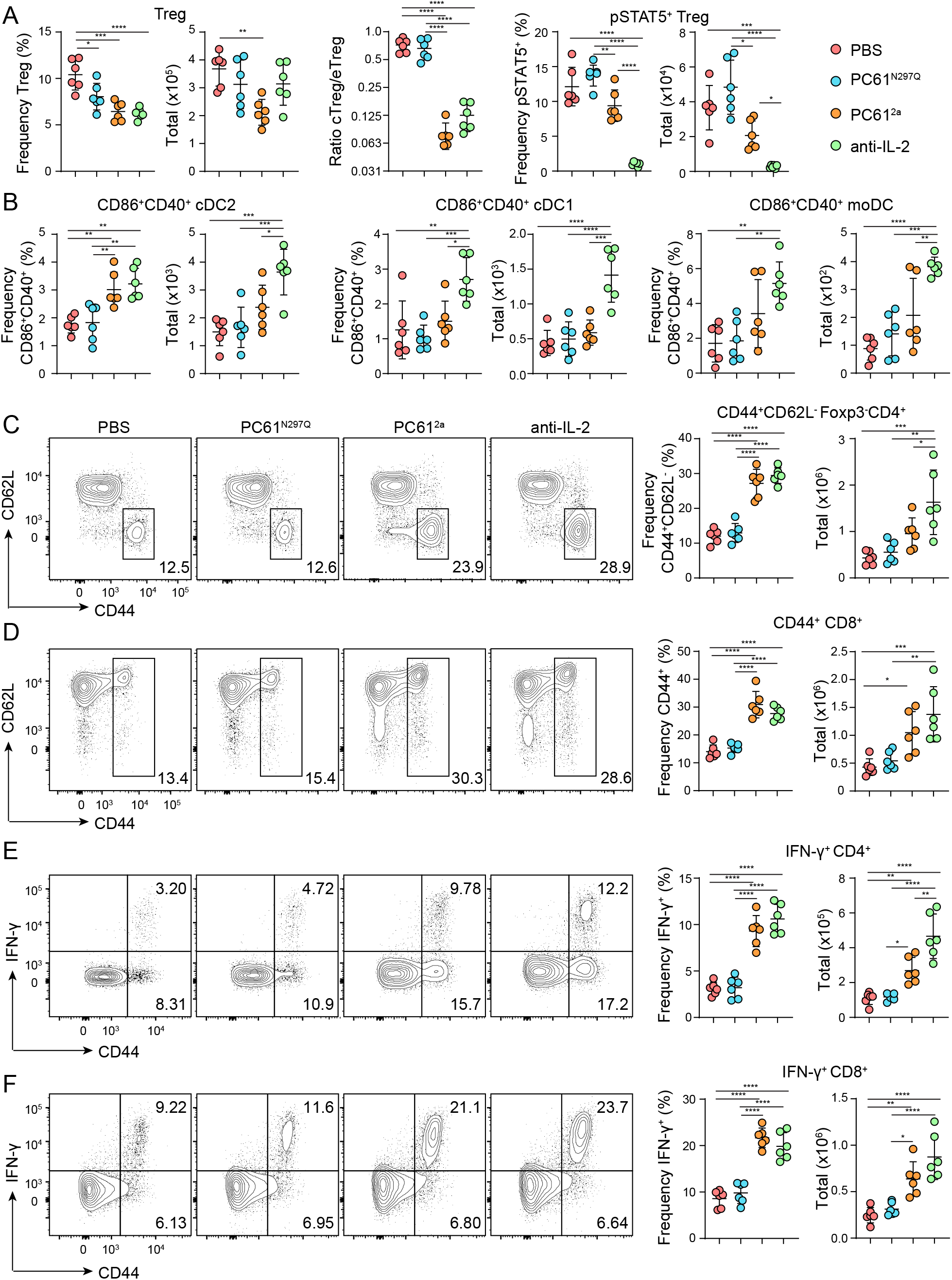
Impact of long-term blockade of IL-2 or CD25 on Treg cells and immune homeostasis. WT B6 mice were treated with PBS, PC61^N297Q^, PC61^2a^, or anti-IL-2 every 7 days, and sacrificed after 28 days for analysis. (A) Left, graphical analyses of frequency and total number of splenic Treg cells in each treatment group. Middle, graphical analyses of the ratio of CD62L^+^ cTreg to CD44^+^ eTreg in the spleens of each treatment group. Right, graphical analysis of frequency and total number of pSTAT5^+^ splenic Treg cells in each treatment group. (B) Graphical analysis of frequency and total number of CD86^+^CD40^+^ cDC2, cDC1, and moDC in spleens of treated mice. Representative analysis of CD44 and CD62L expression by gated splenic CD4^+^Foxp3^-^ T cells (C) and CD8^+^ T cells (D) in each treatment group. Right, corresponding graphical analyses of frequency and total number of gated CD4^+^Foxp3^-^CD44^+^CD62L^-^ (C) and CD8^+^CD44^+^ (D) T cells. (D) Representative analysis of CD44 and IFN-γ expression in gated CD4^+^ (E) and CD8^+^ (F) T cells from each treatment group after stimulation *in vitro* for 4 hours with PMA, ionomycin and monensin. Right, graphical analysis of frequency and total number of IFN-γ producing CD4^+^ and CD8^+^ T cells in each treatment group. Data is combined from two independent experiments, 6 mice per group total. Significance determined by one-way ANOVA with Tukey post-test for pairwise comparisons. *p<0.05, **p<0.01, ***p<0.001, ****p<0.0001.

To examine how *in vivo* treatment with either PC61^N297Q^ or PC61^2a^ affected IL-2 sensitivity, we performed similar dose-response experiments on cells isolated from mice 24h after *in vivo* antibody treatment. Even at this early timepoint, PC61 had saturated all detectable epitopes in mice treated with PC61^N297Q^ or PC61^2a^ (Fig. S2B), and by 7D4 staining we observed reduced CD25 expression in PC61^N297Q^-treated mice, and nearly complete depletion of CD25^hi^ Treg cells in PC61^2a^-treated animals (Fig. 4H). IL-2 sensitivity of CD25^lo^, CD25^mid^ and CD25^hi^ Treg cell populations from treated mice was reduced by about 50-fold (Fig. 4I). However, these Treg cells were still more IL-2 responsive than both CD8^+^ Teff and NK cells (Fig. S2C). Again, further addition of S4B6 to further block IL-2/CD25 interaction severely curtailed IL-2 signaling in treated cells. Together, these data show that although PC61 does substantially reduce the sensitivity of Treg cells to IL-2, sustained CD25 expression and function in PC61^N297Q^ and PC61^2a^ treated mice maintains the Treg cell-dominated hierarchy of access to IL-2 *in vivo*. This continued IL-2 signaling likely underlies the Treg cell function that helps prevent the immune dysregulation observed in anti-IL-2-treated animals.

### Extended treatment with PC61^2a^ disrupts Treg cell-dependent immune homeostasis

While immune dysregulation was only apparent in the anti-IL-2 treated mice after one week of treatment, we wondered if long-term inhibition of CD25 with the PC61^N297Q^ or PC61^2a^ antibodies would ultimately result in loss of Treg cell function. As we observed after one week, the frequency of Treg cells was significantly decreased in mice treated with PC61^2a^ or anti-IL-2 for four weeks, and this predominantly impacted cTreg cells (Fig. 5A). Interestingly, prolonged treatment also resulted in a small but significant decrease in Treg cell frequency in PC61^N297Q^ treated mice (Fig. 5A). However, absolute numbers of Treg cells in the spleen were diminished only in the PC61^2a^-treated mice. Endogenous STAT5 phosphorylation in Treg cells was strikingly similar at four weeks compared to one week, but we now observed activation of cDC2 in both PC61^2a^ and anti-IL-2 treated animals, and enhanced activation of cDC1 and moDC in anti-IL-2-treated mice (Fig. 5B). Accordingly, we detected increased frequencies and numbers of CD44^hi^CD62L^lo^ CD4^+^ (Fig. 5C) and CD44^hi^ CD8^+^ (Fig. 5D) Teff cells in PC61^2a^ and anti-IL-2 treated mice, and this was associated with enhanced production of the pro-inflammatory cytokine IFN-γ by both CD4^+^ and CD8^+^ T cells (Fig. 5E, F). Therefore, continued IL-2 signaling after Treg cell depletion with PC61^2a^ can preserve immune homeostasis only in the short term, while even extended treatment with PC61^N297Q^ does not lead to immune dysregulation.

## Discussion

The importance of CD25 for Treg cell development, survival and suppressive function is well established. Although genetic models using conditional ablation of CD25 on Treg cells demonstrate this [42, 43], it is less clear how therapeutically inhibiting CD25 or IL-2 impacts Treg cells. As the IL-2/CD25 axis is central to mediating immune homeostasis, a precise understanding of how it could be targeted to treat human disease is critical. In this study, we define a progressive cascade of immune dysregulation that occurs upon inhibition of the IL-2/CD25 axis. Full IL-2 blockade results in both a numerical reduction of Treg cells and loss of IL-2-dependent Treg cell functions, thereby leading to rapid immune dysregulation. By contrast, although Treg cell depletion is numerically similar following PC61^2a^ treatment, immune dysregulation is substantially delayed, likely due to the continued IL-2 signaling that supports the function of the remaining Treg cells and potentially aided by depletion of CD25^+^ Teff cells. Finally, although PC61^N297Q^ reduces IL-2 sensitivity ~10-50-fold, this is not sufficient to upset their competitive advantage over Teff cells and NK cells in accessing IL-2, and this has little impact on immune regulation and Treg cell homeostasis even after prolonged treatment.

Whereas previous studies have struggled to distinguish requirements for IL-2 in earlier developmental stages versus subsequent maintenance in adult peripheral immune tissue, we demonstrate here that continued IL-2 signaling in the periphery is critical to maintain Treg cell function. Although maintenance of eTreg cells in non-lymphoid tissues can be IL-2 independent [34, 44], immune steady state in SLOs is rapidly disrupted when IL-2 is neutralized. DCs and Treg cells are linked in a homeostatic loop [45], and work from our lab previously determined that the frequency and function of IL-2 dependent Treg cells in the spleen depends on antigen presentation to autoreactive CD4^+^ T cells largely by CD80/86-bearing cDC2s [9]. Here, we extend these findings to show that when IL-2 is neutralized, costimulatory markers on cDC2s are rapidly upregulated due to reduced functional capacity of the Treg cells in the absence of IL-2 signaling. These data mirror results from Chinen and colleagues [42], in which Treg cells with constitutively active pSTAT5 had an enhanced ability to form conjugates with DCs, resulting in their decreased expression of costimulatory molecules. Treg cells can outcompete naïve T cells to bind to DCs [46], and modulate levels of costimulatory molecules on those DCs through provision of the inhibitory receptor CTLA-4 [47, 48]. The higher Treg cell CTLA-4 expression we observed in the PC61^2a^-treated mice, where Treg cells are able to prevent upregulation of costimulatory molecules on DCs and delay immune dysregulation in contrast to anti-IL-2-treated mice, is likely due to sustained IL-2 signaling. IL-2-dependent regulation of other adhesion and inhibitory molecules could also be involved in providing Treg cells’ enhanced ability to interact with and even strip MHCII-peptide from DCs [49] in order to limit priming of conventional T cells and maintain immune homeostasis.

We further show that Treg cells maintain substantial CD25 function and selective access to IL-2 in a CD25-dependent manner in the presence of PC61, critically clarifying the effects of this commonly used antibody in murine models. Our data strongly support the conclusion that any effects observed in mice treated with PC61 must be due to active depletion of CD25^hi^ Treg cells, and conclusions made in previous studies based on the ability of PC61 to inhibit CD25 function on Treg cells should be re-evaluated [31, 32]. Vargas and colleagues recently demonstrated that tumor-bearing mice treated with a strong depleting CD25 antibody have a significant reduction of intra-tumoral Treg cells and subsequent improved control of tumors, while a non-depleting CD25 antibody has no effect [50]. In cancer therapy this result is desirable. In contrast, more subtle inhibition of the IL-2/CD25 axis, such as treatment with PC61^N297Q^, may be more advantageous for treatment of autoimmunity. For instance, we found that continued IL-2 signaling in splenocytes treated with PC61 was skewed towards the highest CD25 expressing Treg cells compared to the untreated controls. This could be beneficial in an autoimmune setting, particularly in humans where CD25 is expressed on CD56^bright^ NK cells and activated T cells, but at a lower level than on Treg cells [51]. The presence of PC61^N297Q^ could therefore allow a further advantage for Treg cells to preferentially access IL-2 over other effector cells, and improve tolerance in a mechanism similar to Treg selective IL-2 ‘muteins’ [52, 53].

In humans, the anti-CD25 antibodies daclizumab and basilixumab have been used as anti-inflammatory agents to treat MS and prevent graft rejection. This is somewhat counterintuitive given the central role of IL-2/CD25 in maintaining immune tolerance, and the mechanisms of action of these drugs is not well understood. Unlike PC61, these antibodies directly bind to and occlude the IL-2 binding site of CD25, and this results in a reduction in Treg cell frequency (although these cells retain function [54, 55]), increased serum levels of IL-2, and an IL-2-dependent increase in CD56^bright^ NK cells [56–59]. The increase in CD56^bright^ NK cells correlates positively with therapeutic response in patients with MS. While normally thought to be regulatory and more immature, CD56^bright^ NK cells expanded in daclizumab treated patients display enhanced expression of activation markers and receptors that mediate NK cell activation and cytolytic capacity [60], and in vitro studies indicate the ability of these NK cells to kill activated and perhaps encephalitogenic T cells [57, 61]. This increase in bioavailable IL-2 presumably would not occur in patients treated with a CD25 antibody that allowed Tregs to maintain normal levels of signaling, like PC61^N297Q^. While NK cells appear to have a beneficial therapeutic effect, it is also possible they are contributing to adverse events in these treated patients, such as dermatitis, malignancies, infections, and encephalitis [26, 62]. Identification of antibodies that limit CD25 function but allow Treg cells to maintain their selective access to IL-2 may also be therapeutically beneficial in autoimmunity for limiting Teff cell and NK cell responses while maintaining robust Treg cell function.

Targeting of the IL-2/CD25 axis holds incredible promise for treatment of immune dysfunction. Our study emphasizes the complexity of this pathway, and that changes in the sensitivity of cells to IL-2 can produce strikingly different effects on the immune system. Total blockade of IL-2 results in rapid immune dysregulation, while residual CD25 function, even in the face of inhibitory or depleting antibodies, can maintain Treg cell preferential access to IL-2 and therefore allow preservation of immune homeostasis to varying degrees. Subtle differences in CD25 antibody specificity could result in a wide range of beneficial outcomes, and could therefore be utilized to maximize therapeutic benefit.

## Materials and Methods

### Mice

C57BL/6 (B6) mice were purchased from The Jackson Laboratory. DO11.10/Rag2^-/-^ mice were provided by S. Ziegler (Benaroya Research Institute), and soluble OVA (sOVA) mice were provided by A. Abbas (University of California, San Francisco). All mice were bred and maintained at Benaroya Research Institute, and experiments were pre-approved by the Office of Animal Care and Use Committee of Benaroya Research Institute. Mice used in experiments were between 6-20 weeks of age at time of sacrifice.

### Flow cytometry

For DC isolations, minced whole spleens were digested in basal RPMI supplemented with 2.5 mg/mL Collagenase D for 20 minutes under agitation at 37°C. Cell suspensions were then passed through 70 μm strainers into RPMI + 10% FBS (RPMI-10). Erythrocytes were lysed in ACK lysis buffer, and the remaining cells were washed in RPMI-10. DCs were enriched using CD11c-microbeads (Miltenyi) according to the manufacturer’s protocol. Cell surface staining for flow cytometry was performed in FACS buffer (PBS-2% BCS) using the following antibody clones: LiveDead, CD4 (GK1.5, RM4-5), CD8 (53-6.7), CD25 (PC61, 7D4), ICOS (C398.4A), CD44 (IM7), CD62L (MEL-14), NK1.1 (PK136), CD49b (DX5), CD122 (5H4), CD132 (TUGm2), CD5 (53-7.3), CD19 (6D5), Gr-1 (RB6-8C5), CD11b (M1/70), CD11c (N418), MHCII (M5/114.15.2), DC marker (33D1), CD80 (16.10A1), CD86 (GL-1), CD40 (3/23), DO11.10 TCR (KJ1-26), and CD45.2 (104). Cells were incubated in the antibody mixture for 20 min at 4°C and then washed in FACS buffer before collecting events on an LSRII. For intracellular staining, surface antigens were stained before fixation and permeabilization with FixPerm buffer (eBioscience). Cells were washed and stained with antibodies to Foxp3 (FJK-16s), Ki67 (11F6), pSTAT5 (47/pStat5[pY694]), IFN-γ (XMG1.2), and CTLA-4 (UC10-4F10-11). Flow cytometry data was analyzed using FlowJo software.

### Ex vivo staining

To assess pSTAT5 levels directly ex vivo, spleens were immediately disrupted between glass slides into eBioscience FixPerm. Cells were incubated for 20 min at room temperature, washed in FACS buffer, resuspended in 500 μL 90% methanol (MeOH), and incubated on ice for at least 30 minutes. Cells were stained with surface and intracellular antigens, including pSTAT5 (pY694) for 45 minutes at room temperature.

### In vitro assays

For *in vitro* CD25 blockade, splenocytes were isolated from untreated B6 mice as described. 5×10^5^ cells were plated per well into a 96-well round bottom plate. Commercially available PC61 (BioXcell) was added to designated wells at 1μg/mL final concentration, and samples were incubated at 37°C for 30 min and then washed. Meanwhile, 1000 U/mL recombinant IL-2 (eBioscience) was incubated with 50 μg/mL S4B6-1 (BioXcell) for 30 minutes at room temperature. rIL-2:S4B6 complexes were then serially diluted 10-fold to achieve all desired concentrations for the experiment. rIL-2 without S4B6 was subject to the same treatment. rIL-2 or rIL-2:S4B6 dilutions were then added to appropriate wells and samples were incubated at 37°C for 30 min. Samples were then washed and fixed with FixPerm (eBioscience) for 20 min at room temp, washed and incubated in 500 μL MeOH on ice for at least 30 min, washed and finally stained with antibodies for 45 min at room temp. For *in vivo* CD25 blockade, animals were injected intraperitoneally as described with 500 μg PC61^N297Q^ or PC61^2a^. Spleens were harvested 24 hours after injection, and *in vitro* response to IL-2 was measured as described above (without any incubation with commercial PC61).

### Adoptive transfers

Spleen and lymph nodes were harvested from DO11.10/Rag2^-/-^ mice, mashed through a 70 μm strainer and ACK lysed as described above. CD4^+^ (transgenic) cells were enriched by negative isolation according to the manufacturer’s instructions (Dynal). Cells were then enumerated and labelled with CFSE. Finally, labelled cells were washed in PBS before transfer into recipient mice retro-orbitally, 0.5×10^6^ cells per mouse.

### In vivo antibody treatments

For CD25 blocking or depleting, mice were given 500 μg PC61^N297Q^ or PC61^2a^ by intraperitoneal injection every 7 days, or as otherwise specified. For IL-2 blocking experiments, mice were given 150 μg S4B6-1 and either 150 or 500 μg JES6-1A together by intraperitoneal injection every 5 days, or as otherwise specified.

## Acknowledgments

We thank J. Fontenot and Biogen, Inc. for providing engineered PC61 antibodies. We thank S. Zeigler and A. Abbas for providing DO11.10/Rag2^-/-^ mice and sOVA mice, respectively. We thank A. Wojno, K. Arumuganathan and T. Nguyen for help with flow cytometry, and members of the Campbell laboratory for helpful discussions. This work was supported by grants from the NIH to DJC (AI136475, AI124693). ETH was supported by the University of Washington Cell and Molecular Biology Training Grant (5T32GM007270-43).

---

**Figure S1.**
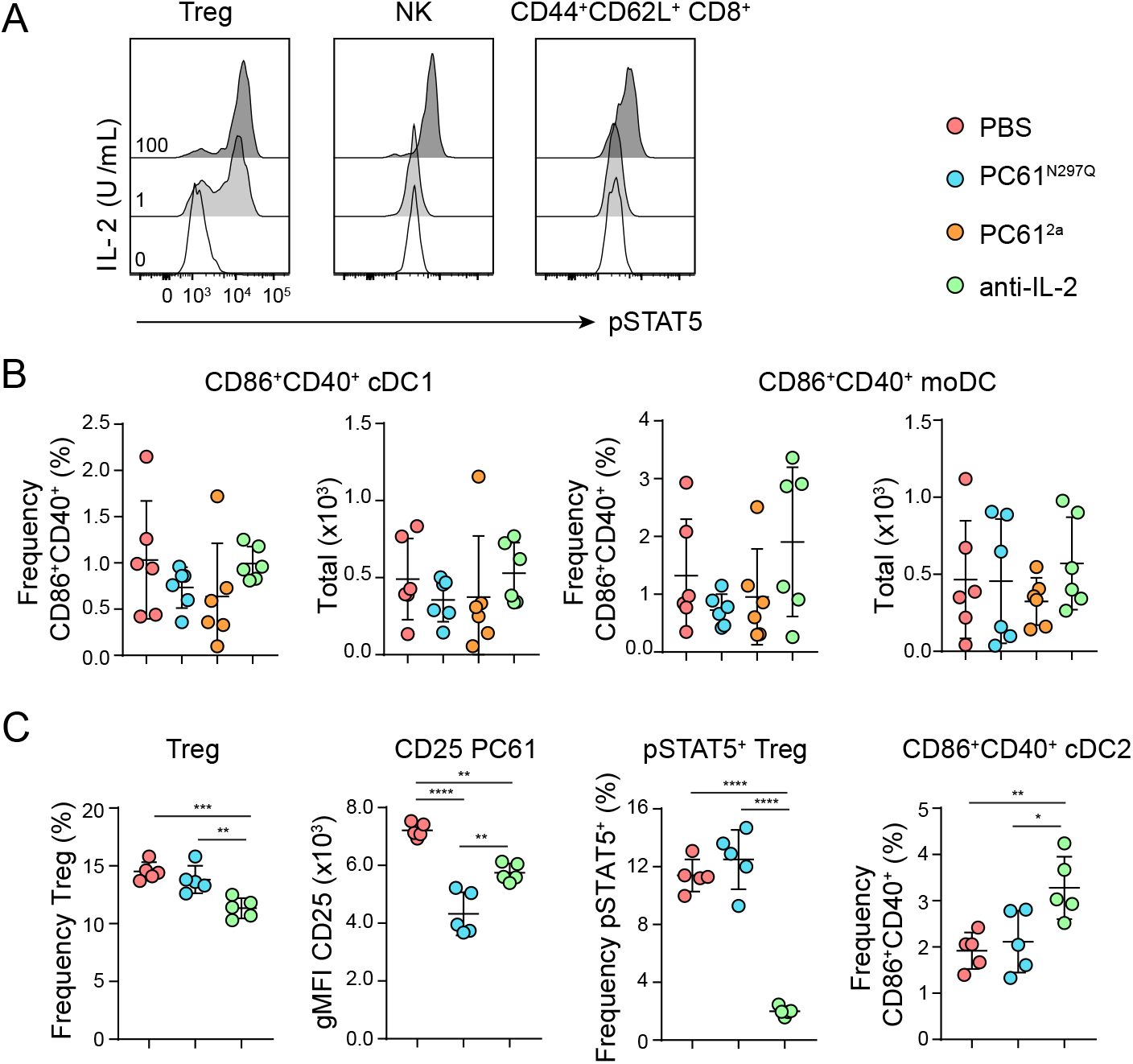
(A) Representative flow cytometric analysis of pSTAT5 in gated splenic Foxp3^+^ Treg cells, NK cells, and CD44^+^CD62L^+^ CD8^+^ cells from WT B6 mice after stimulation *in vitro* with rIL-2. (B) WT B6 mice were treated with PBS, PC61^N297Q^, PC61^2a^, or anti-IL-2 (S4B6 + JES6), and sacrificed for analysis after seven days. Graphical analyses of the frequency and total number of CD86^+^CD40^+^ splenic cDC1 and moDC from all treatment groups. (C) sOVA mice were treated intraperitoneally with PBS, PC61^N297Q^ or anti-IL-2. After seven days, 0.5×10^6^ DO11.10 Rag2^-/-^ CD4^+^ T cells were transferred retro-orbitally into each mouse, and animals were sacrificed five days later for analysis. Graphical analyses of host cell frequency of Treg cells, gMFI of CD25 by Treg cells, frequency of pSTAT5^+^ Treg cells, and frequency of CD86^+^CD40^+^ cDC2 in the spleen of each treatment group. Data is combined from two independent experiments, 5-6 mice per group total. Significance determined by one-way ANOVA with Tukey post-test for pairwise comparisons. *p<0.05, **p<0.01, ***p<0.001, ****p<0.0001.

**Figure S2.**
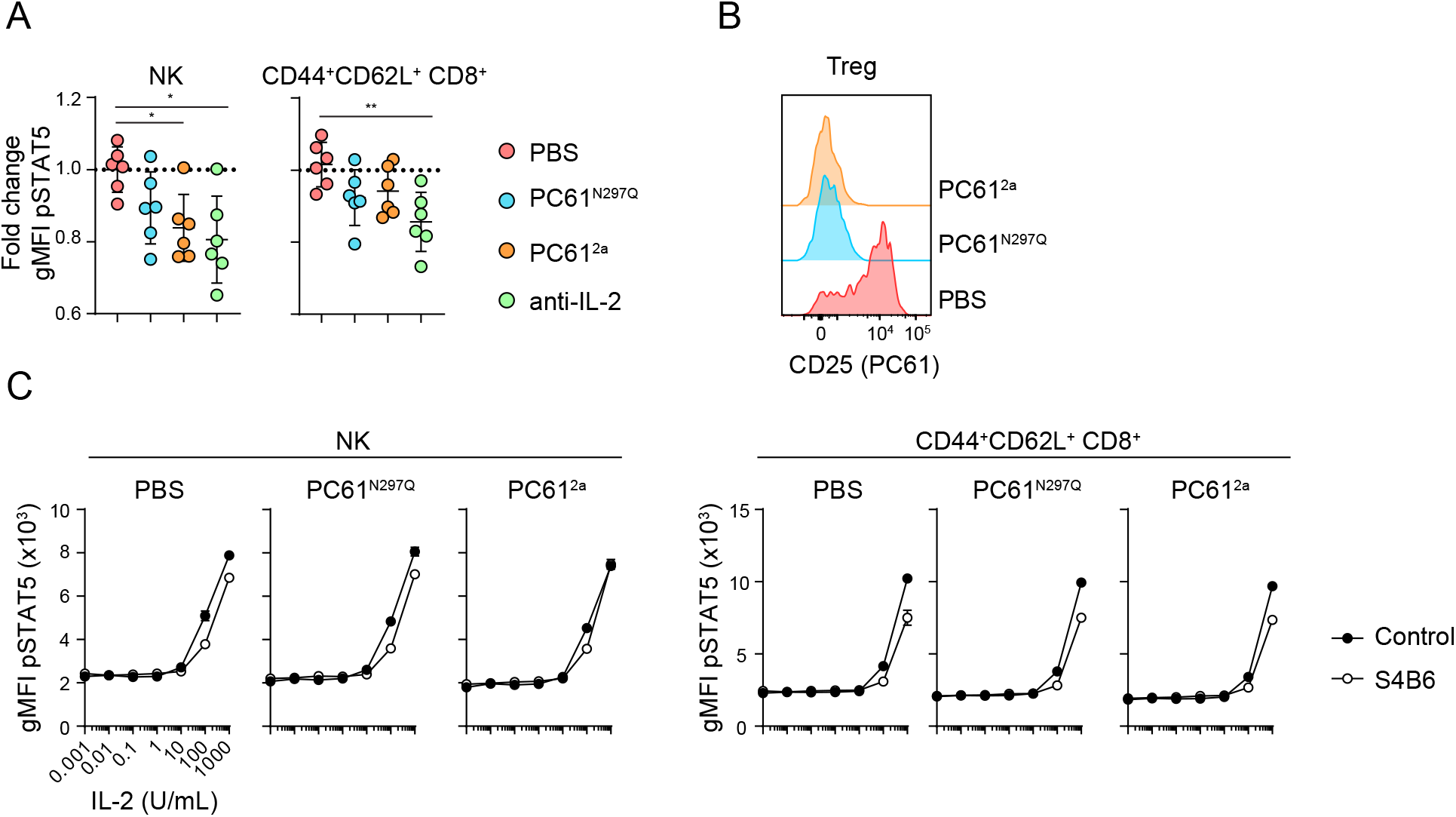
(A) WT B6 mice were treated IP with PBS, PC61^N297Q^, PC61^2a^, or anti-IL-2 (S4B6 + JES6), and sacrificed after seven days for analysis. (A) Graphical analysis of fold change in gMFI over controls of pSTAT5 in NK and CD44^+^CD62L^+^ CD8^+^ cells in each treatment group. Data is combined from two independent experiments, 6 mice per group total. Significance determined by one-way ANOVA with Tukey post-test for pairwise comparisons. *p<0.05, **p<0.01, ***p<0.001, ****p<0.0001. (B-C) WT B6 mice were treated IP with PBS, PC61^N297Q^ or PC61^2a^ and sacrificed after 24 hours. (B) Representative flow cytometry histograms of CD25 (PC61) staining on Treg cells. (C) Graphical analysis of gMFI pSTAT5 NK or CD44^+^CD62L^+^ CD8^+^ cells in response to *in vitro* rIL-2 +/-S4B6 in each *in vivo* treatment group (PBS, PC61^N297Q^ or PC61^2a^). Data is from one representative experiment, with three technical replicates per condition. Experiments were repeated independently at least three times.

